# A microcalorimetric approach for investigating stoichiometric constraints on the standard metabolic rate of a small invertebrate

**DOI:** 10.1101/279562

**Authors:** Ruiz Thomas, Bec Alexandre, Danger Michael, Koussoroplis Apostolos-Manuel, Aguer Jean-Pierre, Morel Jean-Pierre, Morel-Desrosiers Nicole

**Affiliations:** Université Clermont Auvergne, CNRS, LMGE, F-63000 Clermont-Ferrand, France; Université de Lorraine, CNRS, LIEC, F-57000 Metz, France

**Keywords:** Ecological stoichiometry, homeostasis, metabolic rate, *Daphnia magna*, calorimetry

## Abstract

1: Understanding the determinant of metabolism is a core ecological topic since it permits to link individuals energetic requirements to the ecology of communities and ecosystems. Yet, besides temperature, the effects of environmental factors on metabolism remain poorly understood. For example, dietary stoichiometric constraints have been hypothesized to increase maintenance metabolism of small invertebrates, yet experimental support remains scarce.

2: Here, we used microcalorimetric heat flow measurements to determine the standard metabolic rate (SMR) of *Daphnia magna* throughout its ontogeny when fed stoichiometrically balanced (C/P ratio:166) or imbalanced (C/P ratio:1439) diets.

3: When fed a stoichiometrically imbalanced diet, *daphnids* were able to maintain the stoichiometric homeostasis within narrow boundaries. However, they consistently increased their SMR while decreasing their somatic growth rate. Our measurements unequivocally demonstrate that homeostatic regulation implies higher metabolic costs and thereby reduces the portion of energy that can be allocated to growth.

4: Our study demonstrates that microcalorimetry is a powerful and precise tool for measuring the metabolic rate of small-sized organisms and opens promising perspectives for understanding how environmental factors, such as nutritional constraints, affect organismal metabolism.

## Introduction

Metabolism fuels all physiological processes in organisms and appears as the most fundamental mechanism reflecting the energetic cost of living (Hulbert & Else 2000). Individual metabolic rate sets the “pace of life” of a population through its generation time and maximal growth rate (Brown, Gillooly, Allen, Savage, & West 2004; Savage *et al.*, 2004). Therefore, understanding how environmental factors constrain individual metabolic rate, and how these individual-level constraints influence population dynamics of the whole interacting community, is key for understanding ecosystems (Pawar, Dell, & Savage 2015). Standard metabolic rate (SMR) represents the minimum energy that ectotherms must expend on tissue maintenance but also on homeostatic mechanisms needed to sustain life (Fry 1971, Auer, Salin, Rudolf, Anderson, & Metcalfe 2015). Homeostasis capacity of organisms is at the core of ecological stoichiometry (ES) approach which stipulates that the imbalance of elements, such as carbon, nitrogen or phosphorus between consumers requirements and nutritional resources, force consumers to activate specific regulatory processes. Ecological stoichiometry posits that such processes are energetically costly and may thereby affect SMR (Hessen, Elser, Sterner, & Urabe 2013).

A large number of studies have been devoted to investigate the effects of nutrient limitation on invertebrate performance (Elser *et al.*, 2000; Sterner & Elser 2002; Frost, Stelzer, Lamberti, & Elser 2002; Cross, Benstead, Frost, & Thomas 2005; Hessen *et al.*, 2013), yet only a few have been dedicated to elucidate physiological processes involved in maintaining homeostasis and even less considered effects on metabolic rate. However, in 1994, Sterner and Hessen already hypothesized that consumers could cope with a dietary C excess by increasing their respiration rate. This hypothesis is well accepted despite the fact that only a handful of studies attempted to unravel it experimentally (Zannoto, Gouveia, Simpson, & Calder 1997; Darchambeau, Faerovig, & Hessen 2003; Jensen and Hessen 2007; Jeyasingh 2007). The main difficulties for testing this hypothesis rely on the low sensitivity and the numerous biases related to respirometric methods (Walsberg & Hoffman 2005). These issues are further exacerbated when considering relatively small animals such as insects or zooplankton species, often used as models in nutritional ecology but generating very low O_2_ and CO_2_ fluxes. Recent technical and methodological improvements (Rosewarne, Wilson, & Svendsen 2016) might help to reduce these biases to some extent, yet a careful interpretation of the results is needed (Clark, Sandblom, & Jutfelt 2013). More recently, Lukas and Wacker (2014) and Elser *et al.* (2016) could confirm that mass-specific respiration rates increased when organisms consumed imbalanced diets. However, both of these studies were performed on feeding animals and did not attempt to discriminate the specific dynamic action (SDA, i.e. the increase in respiration associated with feeding) from standard metabolic rate. SDA can be highly variable and represent up to 30% of total metabolic rate (McCue 2006) and thereby has to be excluded in order to determine SMR. Finally, despite the increased precision of respirometric equipments, the translation of respirometric measurements into metabolic energy equivalents still relies on a set of strong assumptions (Walsberg & Hoffman 2005). The latter include: (1) the nature of the primary substrate that is catabolized (i.e. lipids, proteins or carbohydrates), (2) its degree of oxidation and (3) the absence of O_2_-consuming or CO_2_- producing synthetic processes. All of these assumptions can be violated depending on the nutritional status of the assayed individuals (Walsberg & Wolf 1995; Walsberg & Hoffman 2005) thereby hindering up to now a clear assessment of the relationship between dietary stoichiometric constraints and SMR.

From a methodological perspective, calorimetry constitutes the “*golden standard*” for quantifying metabolic rates (Kaiyala & Ramsay 2011). It allows the direct and non-invasive assessment of the metabolic energy produced by an organism measured in terms of heat dissipated by all chemical reactions and energy transformations taking place in the cells. Considering this, calorimetry should enable a precise determination of organismal metabolic rate. Quite surprisingly, despite its early success (for history of animal calorimetry see Kleiber 1961 and Lamprecht & Schmolz 1999) and high potential, calorimetry remains largely underused in more recent ecological research (but examples of its use can be found in the minireview of Braissant, Wirz, Gopfer, & Daniels 2010, in Bricheux, Bonnet, Bohatier, Morel & Morel-Desrosiers 2013 and in Lescure, Carpentier, Battaglia-Brunet & Morel-Desrosiers 2013) and even less in animal ecology (but see Normant, Becker & Winkelmann 2015). This might be explained by the lower sensitivity and complexity of older instruments that required working with organisms large enough to produce measurable heat flows (Normant, Dziekonski, Drzazgowski, & Lamprecht 2007) but also by the increased availability of affordable and user-friendly respirometers (Kaiyala & Ramsay 2011). Fortunately, modern microcalorimeters, which have become extremely sensitive and easier to use, now enable direct measurement of the heat produced by organisms as small as tiny insects or zooplankton species.

Here, we experimentally illustrate the high potential of microcalorimetry in ecological studies by determining the metabolic response of homeostatic consumers undergoing a dietary stoichiometric constraint. We hypothesize that the necessary maintenance of homeostasis of consumers induces a higher SMR in organisms fed a stoichiometrically imbalanced diet. To test this hypothesis, we compare, in a standardized growth experiment, the individual heat flow produced versus mass of a clonal line of *Daphnia magna* submitted or not to a stoichiometric constraint during its ontogenetic growth.

## Material and Methods

### Origin and maintenance of daphnids

The planktonic cladoceran *Daphnia magna* used in this study is a classical model organism in ecological stoichiometry approaches (Hessen et al., 2013) and biology in general (Altshuler *et al.*, 2011) due to its short development cycle and parthenogenetic reproduction allowing to work with clonal population. *Daphnia* is a non-selective filter feeder preponderant in freshwater communities and considered as a keystone species in planktonic food webs (Dodson & Hanazato 1995). The *Daphnia magna* clone used here was collected from a pond of the Allier River floodplain (Puy-de-Dôme, France). Females were maintained for several generations in laboratory in 2L glass container of Volvic© water on a 14:10 h light:dark cycle at 20°C and with a maximum of 15 individuals per litre. They were fed daily *ad libitum* (3mg C.L^−1^ well above the incipient limiting level that is reported to be approximately 0.5mg C.L^−1^; Lampert 1978) with a suspension of the green algae *Chlamydomonas reinhardtii.*

### Algae cultures and preparation

*Chlamydomonas reinhardtii* was used as food during all the experiment. Algae were grown on modified WC medium with vitamins (Von Elert & Wolffrom 2001) with 7.1 mg.L-1 or 0.35 mg.L-1 phosphates (PO_4_) providing respectively a control algae (P+) or a P-deficient algae (P-). Cultures were maintained at 20°C under permanent light at a dilution rate of 0.10 per day using aerated 5L vessels. Food suspensions were daily prepared from stock solution of algae by centrifugation and resuspension of the cultured cells in Volvic© water. The carbon concentrations of the algal food suspensions were estimated from photometric light extinction (800 nm) and pre-established carbon concentration - light extinction regressions. Algal culture subsamples were taken at the beginning, the middle and the end of the experiment, filtered and dried to measure their carbon and phosphorus content.

### Experimental setup

Before the experiment, one individual was isolated from the stock culture of *Daphnia magna* and maintained in the same conditions as described earlier. When this clone produced offspring, the neonates (first generation) were kept and the mother removed. This step was repeated twice to keep females from the third generation. After they had released their first and second clutch, females from the third generation were kept and the neonates discarded. In order to minimize inter-individual variability, neonates from the third clutch and less than 8h-old were used for the experiments

During the first 24h, the daphnids were fed on a suspension of P+ *Chlamydomonas reinhardtii* (3mg C.L^−1^) and then randomly distributed into 4 replicate flasks for each treatment (P+ and P-). In each of these 500mL flasks, organisms were raised in Volvic© water renewed each day and had a daily *ad libitum* food supply (3mg C.L^−1^) to maintain optimal feeding conditions during the 9 days of the experiment. Each day of measurement, one individual was randomly selected from three of the four replicate flasks to determine its metabolic rate by microcalorimetry resulting in triplicate measurements per dietary treatment and per day of measurement (except for day 9 studied in quadruplicate). After measuring their heat flow, the organisms were sacrificed and individually placed into pre-weighed aluminium containers, dried 48h at 60°C and weighed (Sartorius ME36S; Germany) to determine their dry weight. Simultaneously, in the same replicate flasks, a pool of five organisms was removed for somatic growth rate, C and P content measurements.

### Carbon and Phosphorus analysis

After drying 48h at 60°C, filtered algae were crushed and placed with 10mL ultra-pure water in 20 ml vials. Daphnids were dried, reduced to powder and placed with 10mL ultra-pure water in 20 ml vials. The following steps are similar for both daphnid and algal samples: Carbon content was measured directly with a C analyser (Shimadzu: TOC-V; Japan). Phosphorus content was estimated using the potassium persulfate method as describe by Gales, Julian, and Kroner (1966). Briefly, potassium persulfate and heating transform organic phosphorus in 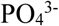 and then a solution of sulfuric acid, ammonium molybdate, ascorbic acid and antimony-potassium tartrate transform 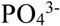 in a blue solution of phosphomolibdic acid. The intensity of the colour is proportional to the quantity of 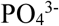 transformed allowing a spectrophotometric measurement of initial P content of the sample. Carbon and phosphorus content are then related to dry weigh to establish C and P proportion in organisms and their molar C/P ratio.

### Growth rate measurement

After 48 h drying, organisms were weighed with a precision of 0.001 mg using a micro-balance (Sartorius ME36S; Germany). They were then placed at 60°C one more day and re-weighed to ensure their total desiccation and a maximum precision of the measurement. At the beginning of the experiment, triplicates of 20 randomly selected neonates were weighed in the same conditions to determine the initial dry weigh.

Growth rate was calculated as follow: **Growth rate = (ln(W_t_) – ln(W_0_)) / t**

where W_t_ is the weight (in μg) at age t (in days) and W_0_ the weight (in μg) of neonates.

### Heat flow measurement by microcalorimetry

Isothermal microcalorimeters of the heat-conduction type are particularly interesting because they allow direct record of the heat flow as a function of time. Here, a TAM III (Waters, TA Instruments; United States) equipped with two multicalorimeters, each holding six independent minicalorimeters, was used to record continuously and simultaneously the heat flow of 12 samples, with a precision of at least ± 0.2μW. The temperature of the bath was set at 20°C and its long-term stability was ± 1×10^−4^ °C over 24 h. The minicalorimeters were electrically calibrated. Specific disposable 4 mL ampoules, capped with butyl rubber stoppers and sealed with aluminium crimps, were used in each minicalorimeter. Daphnids were rinsed three times with Volvic© water to remove all food particles and then were kept one hour in Volvic© water to exclude excretory products. After that, they were placed individually into the microcalorimetric ampoules containing 2 mL of Volvic© water. In preliminary experiments, we verified that oxygen diffusion from the 2 mL of air present in each ampoule was sufficient to ensure aerobic conditions during the whole measurement duration. Three ampoules were prepared for each experimental condition. Some ampoules were also prepared with only water in order to carry out blank experiments. The closed ampoules were introduced into the 12 minicalorimeters and the data collection was simultaneously launched using the TAM Assistant Software (Waters, TA Instruments; United States). A heat flow value was collected every second during at least 8 hours for each of the samples studied. Blank experiments showed that after 2 hours of temperature equilibration the microcalorimetric signal reaches a plateau at zero, confirming that our experimental conditions did not induce any significant bacterial growth (Fig.1). In the presence of daphnids, the experiments showed that the signal still decreases slowly after 2 hours until it reaches a plateau after an interval of about 6 hours. The heat flow value corresponding to the steady-state was thus determined, in all cases, by fitting linearly the signal between the 6^th^ and 8^th^ hours (Fig. 1). With this method we were able to determine precisely the heat flow produced by individuals of different sizes, even those weighing only a few μg (Fig.2). Moreover, we showed that the heat produced by individuals bred in the same conditions but of different sizes were clearly distinguishable and repeatable (Fig. 2).

**Figure 1:**
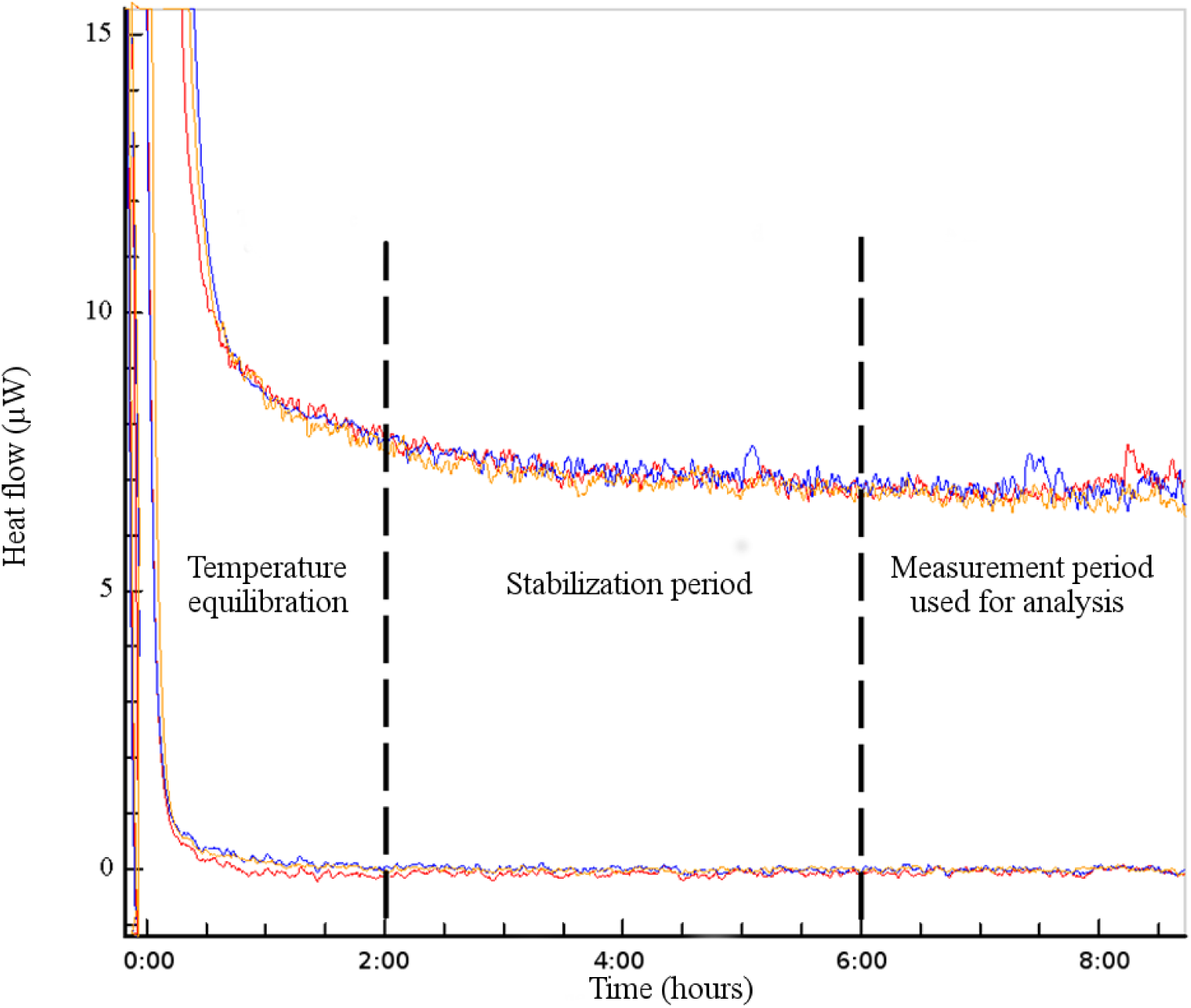
Examples of individual Heat Flow (μW) versus Time (hours) curves: blank experiments (lower lines); experiments with 9-day-old control fasting daphnids (upper lines). Each color represents an individual (n=3)

**Figure 2:**
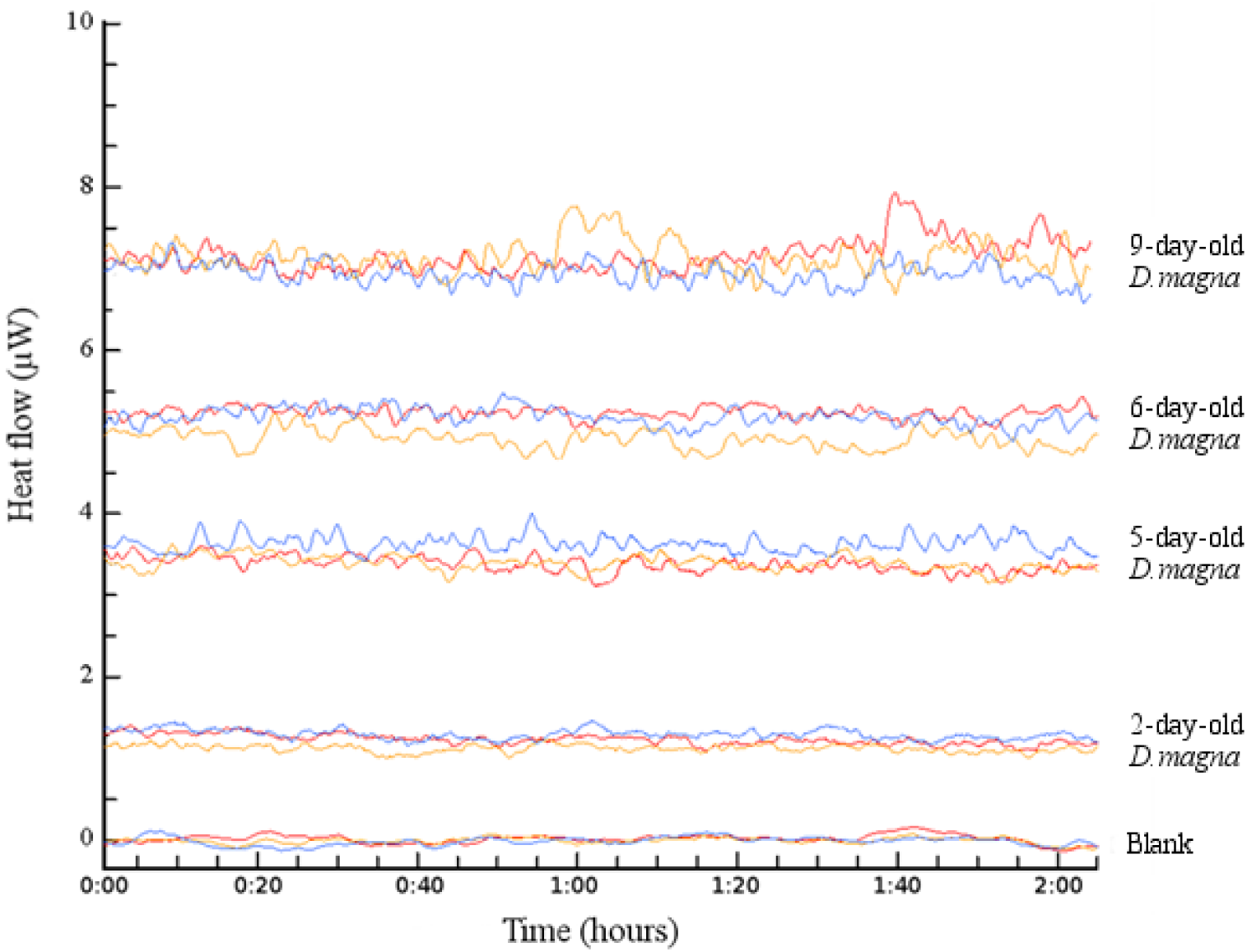
Individual Heat flow (μW) versus Time (hours) curves observed in the absence of daphnids (blank) and in the presence of 2-, 5-, 6- or 9-day-old control fasting daphnids. Each color represents an individual (n=3)

### Data analysis

Prior to analysis, all data were log_10_ transformed in order to meet ANOVA assumptions. Algae and daphnid C and P content as well as C/P ratio of the two dietary treatments were compared for each experimental duration using a two way ANOVA. Differences in daphnid individual metabolic rate (i.e. heat production) between stoichiometically balanced or imbalanced diets were compared by ANCOVA using individual mass as covariate. All statistical analysis were performed using R v.3.4.3 (R Core Team 2018) with an α error set at 0.05.

### Results

## Organisms’ mineral content

The green algae *Chlamydomonas reinhardtii* of the P+ treatment showed a ten-fold lower C/P than P-*C. reinhardtii* (F_(1;4)_=1032.3; p<0.001) but no temporal variability was observed during the experiment (F_(1;4)_=1.88; p>0.1). The carbon relative content was similar in the two treatments (F_(1;4)_=10.73; p>0.05)) and the variation of C/P ratio was only explained by a ten-fold reduction of P in P-algae (F_(1;4)_=1296.9; <0.001) (Table 1).

**Table 1:**
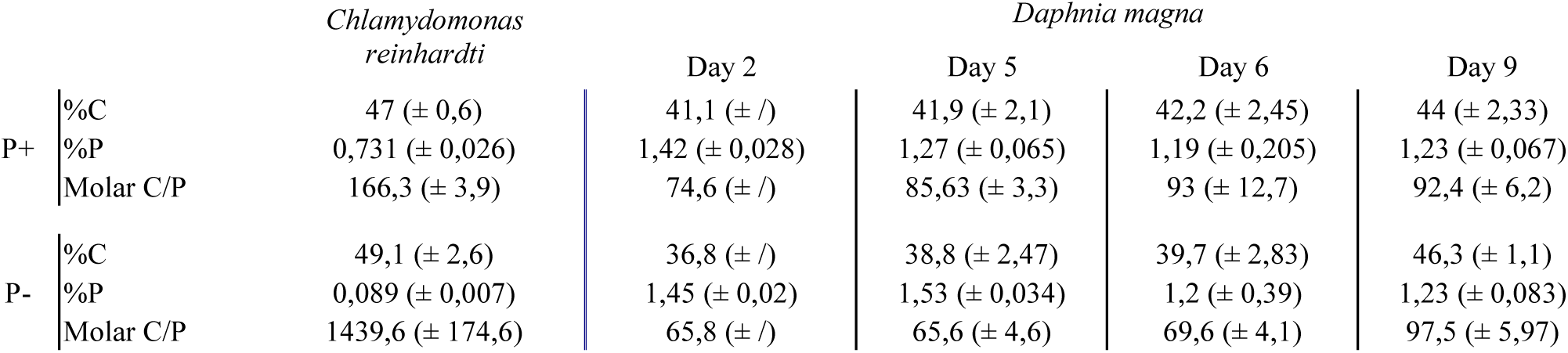
*Mean (± s.e., n=6) Carbon, Phosphorus content and C/P molar ratio of algae (Chlamydomonas reinhardtii) and experimental organisms (D.magna) in the control (P+) or P depleted (P-) treatments. C and P are given in % of total dry mass.*

Treatment (P+ or P-) had no significant effect on the C in daphnids which accounted for ca. 40% of their dry weight (F_(1;20)_=1.63; p>0.1). Daphnids fed P-had a significantly higher proportion of phosphorus during the experiment (F_(1;20)_*=9.37; p<0.01*). Molar C/P ratios of daphnids appeared lower in the P-than in the P+ treatment (F_(1;20)_=17.9; p<0.001). A significant interactive effect between diet treatment and age on the daphnid individual C/P ratio (F_(3;20)_=7.2; p<0.01) was found, explained by a stronger influence of diet C/P on younger individuals.

### Daphnia growth

Organisms from P-treatments exhibited a significantly lower growth rate than those from P+ (Fig. 3) (F_(1;24)_=23.68; p<0.001). Daphnids’ age also induced a significant variation in growth rate (F_(3;24)_=46.82; p<0.001) where younger organisms exhibited a higher growth rate than the older ones (Fig.3). Dietary stoichiometry and organisms age also had an interactive effect on growth (F_(3;24)_=3.94; p<0.05) with younger individuals being more responsive to the dietary C/P ratio.

**Figure 3:**
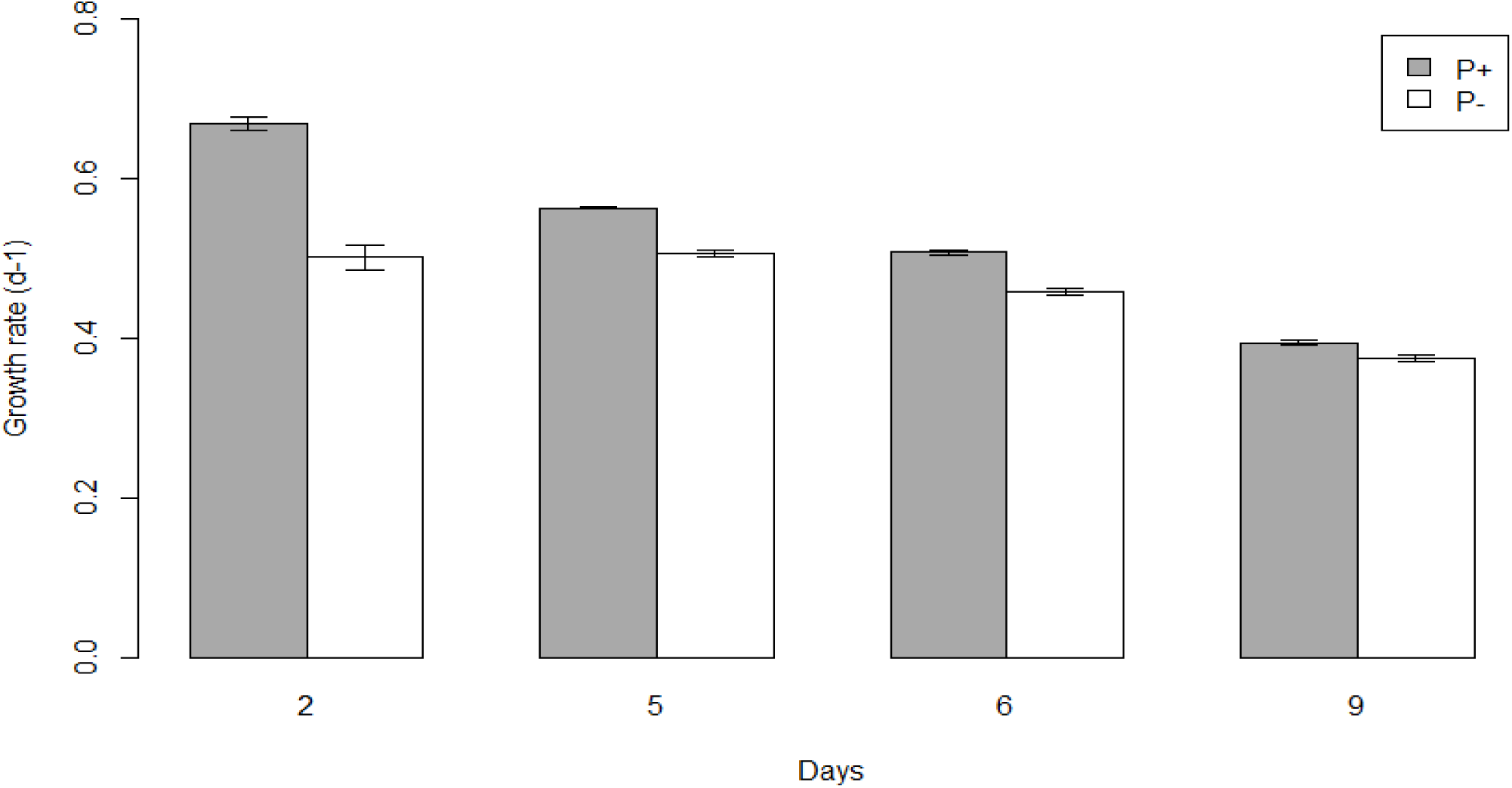
*Mean (± s.e., n=3 except day 9 where n=4) D. magna growth rate (day*^*-^1^*^*) versus age (days) for the control (P+) and the P-depleted (P-) treatments.*

**Figure 4:**
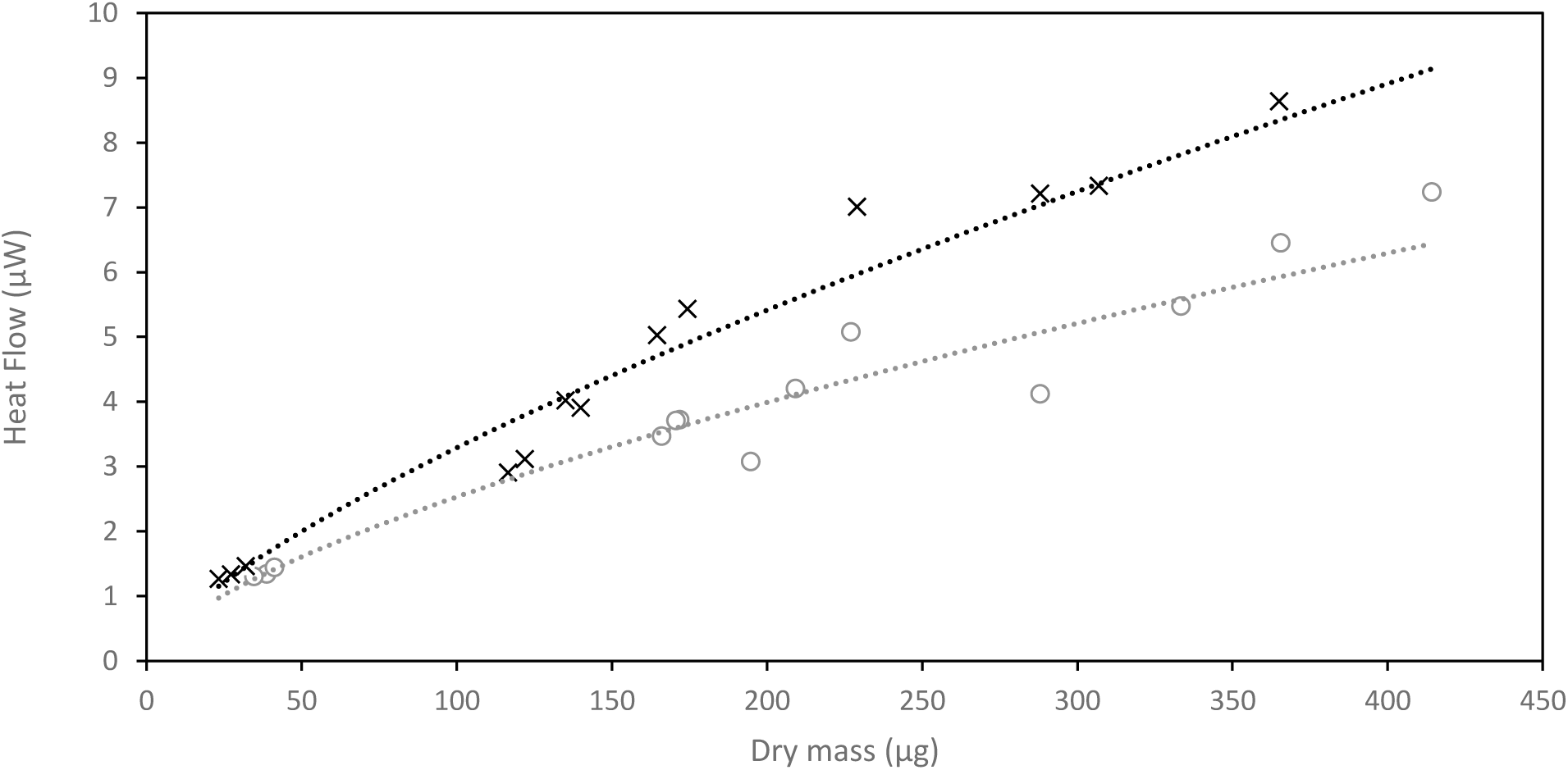
*Individual heat flow (μW) versus body dry mass (μg) for treatment P+ (grey circles; regression: y=0.12x*^*0*^.^*66*^; *R*^*2*^*=0.96) or P-(black cross; regression: y=0.12x*^*0*^.^*72*^; *R*^*2*^*=0.97).*

### Heat flow produced by daphnids

We found a significant effect of the dietary treatment on organisms’ metabolic rate after controlling for individual body mass (F_(1;23)_=36.03; p<0.001) (Fig.4), organisms from the P-treatment showing a higher heat flow that those from the P+ treatment. Individual mass had also induced a significant variation of the heat produced by organisms (F_(1:23)_=616.02; p<0.001). No interactive effect of treatment and mass has been found (F_(1;23)_=1.31; p>0.05) (Table 2).

**Table 2:**
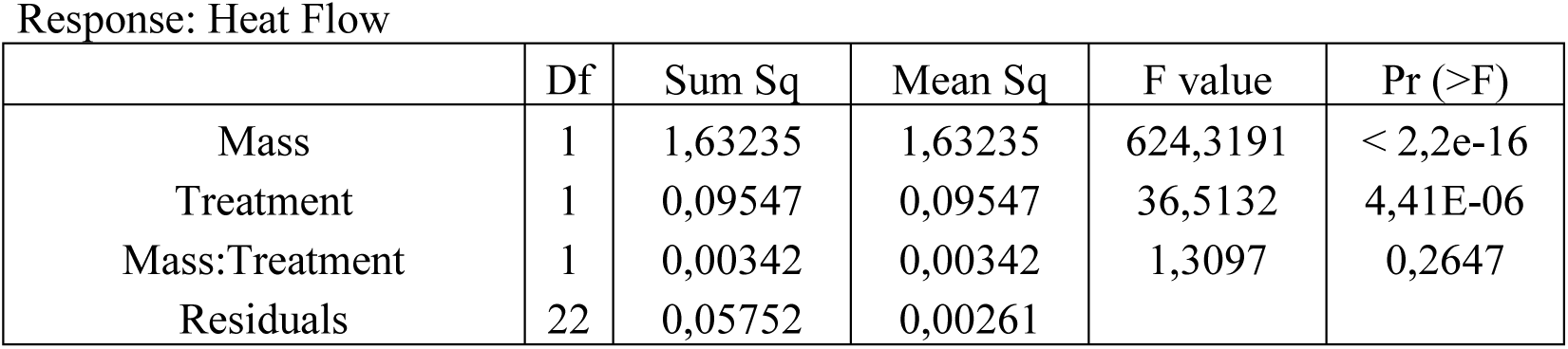
Summary of the ANCOVA analysis performed to determine effects of mass and treatments on individual heat flow.

## Discussion

We conducted a growth experiment on a clone of *D. magna* to evaluate how a dietary stoichiometric constraint could affect its standard metabolic rate (SMR). Isothermal microcalorimetry enabled high precision and repeatability of SMR measurement (Fig. 1-2). We were therefore able to demonstrate unambiguously and with unprecedented precision the age and diet-related changes in the SMR of *D. magna*.

As SMR is the most fundamental component of organismal energy budget (Sibly *et al.*, 2013), its variation should affect the energy that can be allocated to other functions such as growth or reproduction (Burton, Killen, Armstrong, & Metcalfe 2011; Sibly *et al.*, 2013). Most studies on the variation of SMR have focused on inter-genotype differences and their consequences on individual fitness. Some studies suggested that a higher SMR confers a fitness advantage by maintaining larger metabolic machinery facilitating higher resources intake (Metcalfe, Taylor, & Thorpe 1995; Yamamoto, Ueda, & Higashi 1998; Careau *et al.*, 2012). Conversely, others found that the energetic cost of SMR is a burden to the organismal energy budget and that a low SMR should let a greater amount of energy available for growth or reproduction (Alvarez & Nicieza 2005; Blackmer *et al.*, 2005; Reid, Armstrong, & Metcalfe 2011). However, as environmental factors are likely to affect individuals metabolic rate (Glazier *et al*., 2011), interpretingnindividual variation in SMR requires an understanding of the relative importance of environmental factors in comparison to genetic variations of SMR (Glazier 2010, Rosenfeld, Van Leeuwen, Richards, & Allen 2015). Thus, our focus here is not on genotype differences in SMR but on the link between SMR and an environmental factor (diet quality). We therefore used a single *Daphnia* clone to minimise inter-individual genetic differences.

In our experiment, body C/P ratio of *D. magna* showed significant variations depending on both organisms age (size) and dietary stoichiometry. This interactive effect is not surprising since younger individuals of *Daphnia* classically present higher phosphorus contents than adults (DeMott 2003). This observation has been explained by the potentially higher growth rate of juveniles when compared to adults, faster growth leading to higher P-requirements (Growth Rate Hypothesis, Elser *et al.*, 2003). In our experiment, the lower C/P ratio of organisms fed low-P resources could thus be explained by their smaller sizes during the experiment. This result confirms that P requirements can change with organisms ontogeny (Halvorson & Small 2016) but do not question the homeostatic regulation capabilities of daphnids. During our experiment our results clearly showed that, regardless of their size, daphnids facing an imbalanced diet had to regulate their body stoichiometry by releasing excess C. Under certain conditions, such stoichiometric regulation can occur through pre-absorptive pathways, through adjustments in ingestion or assimilation rate (DeMott, Gulati, & Siewertsen 1998; Plath & Boersma 2001). However, previous studies (Darchambeau *et al*, 2003) showed that when fed *ad-libitum* as in our experiment the ingestion or assimilation rates of *Daphnia* are not affected by dietary stoichiometry. Hence, we can assume that the homeostatic regulation we observe is mainly explained by post-absorptive processes. Such processes involve an active release of C, either through a higher excretion of DOC and/or a greater respiratory release of CO_2_ (Jensen & Hessen 2007; Lukas & Wacker 2014) leading to the hypothesis that a dietary stoichiometric constraint increases SMR (Hessen et al 2013).

Standard metabolic rate represents the very minimum energy required for organism’s maintenance corresponding to the metabolic activity of a resting, non-feeding organism at a given temperature (Fry 1971, Auer *et al.*, 2015). However, measuring the exact value of SMR is usually difficult due to some unavoidable organismal routine activity (Willmer, Stone, & Johnston 2005). As an example, pelagic zooplankton species, such as *Daphnia*, must remain continuously active to prevent sedimentation. This swimming activity is constant and essential for survival, we therefore argue that it is consistent to consider it as part of the maintenance processes. On the other hand, specific dynamic action (SDA, see McCue 2006) complicates metabolic data interpretation, and should be excluded at best in order to measure SMR. In this regard, our method, which allows a temporally unconstrained (as opposed to microrespirometry which is constrained by the oxygen level in the microwells) real-time monitoring of the organismal heat flow, presents a clear advantage since it makes it possible to determine precisely when SDA becomes negligible (i.e. heat flow reaches a plateau). It has indeed been shown here that, whereas the microcalorimetric signal of pure water reaches a steady-state relatively rapidly, its decrease reflecting simply the temperature stabilization of the sample, the heat flow produced by the daphnids keeps decreasing for a few hours after the food is removed. We interpret this gradual decrease of the heat flow during the fasting period as the decrease of SDA to a negligible level. This observation provides a useful temporal range for the arrest of digestive activity following food removal in *Daphnia*. By eliminating the various sources of bias discussed above, our experiments demonstrate that daphnids facing a dietary stoichiometric constraint increase their SMR reflecting higher costs of homeostatic body stoichiometry regulation. Under the assumption that the ingestion and assimilation rates remain constant across the applied treatments (Boersma & Wiltshire 2006), this increase of SMR should alter the energy budget by reducing the energy available for growth. This is strongly supported by the significantly decreased growth rates in the treatments with the highest SMR.

In conclusion, it has been shown that isothermal microcalorimetry of the heat-conduction type allows a precise, reproducible, continuous and real-time monitoring of the metabolic rate of small invertebrates. By using a multichannel instrument, several samples can be studied simultaneously (12 in the present case). For each of the studied samples, a heat flow value can be collected every second during several hours, which leads to an outstanding delineation of the heat flow – time curve. This non-stop monitoring allows the exclusion of SDA and thus a precise determination of SMR. In a more general context, the possibility of metabolic activity real-time monitoring without any temporal constraint offers the opportunity to understand metabolic rate variation at different temporal scales ranging from minutes to days. Very low detection limit, real-time monitoring and high-throughput data collection clearly give to microcalorimetry an advantage over more classical techniques for the determination of the metabolic rate and make it a powerful tool for investigating animal metabolism in various ecological contexts.

## Acknowledgments

This study contributes to the **PHOCSAPASS** research project funded by the **EC2CO/ CNRS** program.

